# Computational models for state-dependent traveling waves in hippocampal formation

**DOI:** 10.1101/2023.05.19.541436

**Authors:** Yuxuan Wu, Zhe Sage Chen

## Abstract

Hippocampal theta (4-10 Hz) oscillations have been identified as traveling waves in both rodents and humans. In freely foraging rodents, the theta traveling wave is a planar wave propagating from the dorsal to ventral hippocampus along the septotemporal axis. Motivated from experimental findings, we develop a spiking neural network of excitatory and inhibitory neurons to generate state-dependent hippocampal traveling waves to improve current mechanistic understanding of propagating waves. Model simulations demonstrate the necessary conditions for generating wave propagation and characterize the traveling wave properties with respect to model parameters, running speed and brain state of the animal. Networks with long-range inhibitory connections are more suitable than networks with long-range excitatory connections. We further generalize the spiking neural network to model traveling waves in the medial entorhinal cortex (MEC) and predict that traveling theta waves in the hippocampus and entorhinal cortex are in sink.

## Introduction

The hippocampus is important for episodic memory, navigation, learning and planning. Its subneworks are traditionally viewed as multisynaptic, unidirectional circuit organization (EC→DG, DG→CA3, CA3→CA1) that promotes propagation of excitatory processes (1–3). The hippocampal subneworks are traditionally viewed as having a feed-forward, unidirectional circuit organization that promotes propagation of excitatory processes (3). The septotemporal axis of the hippocampus (a.k.a. dorsoventral or longitudinal axis in the rodent) can be subdivided into dorsal (septal), intermediate, and ventral (temporal) portions based on their distinct entorhinal inputs, subcortical projections and gene expression (4–6). The multisynaptic loop is orthogonal to the dorsoventral axis and is present at all septotemporal levels. Since inputs to the different segments of the hippocampus from other brain areas differ, different functions are attributed to these segments. For example, the dorsal hippocampus is more involved in spatial navigation, whereas the ventral hippocampus plays a prominent role in emotional behavior (5). On the other hand, the transverse (proximal-distal) axis of the hippocampus has also given rise to functional diversity in hippocampal coding (7) and distinct place coding in CA1 (8). Theta oscillations (4-10 Hz), originated in the medial septum, provide the major feedforward inhibition to CA1 or CA3 inhibitory neurons. Theta power decreases from the dorsal to ventral hippocampus, suggesting that inhibition is strongest in the dorsal segment.

Identification and detection of spatiotemporal neural patterns have become possible with the advances in large-scale high-density neural recordings (9–11). Traveling waves are spatiotemporal patterns with steady phase shifts as a function of space, which has been widely reported in the hippocampus (12–15) and other cortical areas (16–23). Specifically, hippocampal traveling waves are state-dependent, appearing in the form of propagating waves of theta oscillations during awake run behavior and rapid-eye-movement (REM) sleep (12,13), or in the form of sharp-wave ripples (SWRs) (120-150 Hz) during rest or non-REM (NREM) sleep (24). Mesoscopic traveling waves may arise from multiple distinct mechanisms, such as a signal pacemaker generator, or chains of unidirectional linked neurons (25), or a network of weakly coupled oscillators (26). Several functional roles of hippocampal or cortical traveling waves have been suggested (17), but a complete understanding of the computational mechanism remains elusive. Several computational models have been developed for explaining various cortical traveling waves, including the Wilson-Cowan model (27), coupled oscillators (25,28), neural field model (29,30), Izhikevich model (31,32), and large-scale circuit modeling based on leaky integrate-and-fire neurons (LIFs) (33,34). However, computational models for the propagation of hippocampal theta waves are relatively unexplored (35).

In this paper, we develop biologically inspired large-scale computational models consisting of excitatory and inhibitory LIF spiking neurons, which are motivated by the current knowledge of rodent hippocampal anatomic connectivity and physiology. By computer simulations, we describe conditions under which hippocampal traveling waves can be generated and propagated. Our computational models accommodate two distinct assumptions of long-range synaptic connectivity based on either excitatory or inhibitory connections. We characterize the traveling wave features with respect to running speed and brain states, and further generalize the model to theta traveling waves in the medial entorhinal cortex (MEC) (36). Our models also provide experimentally testable predictions and new insight into detection of planar traveling wave patterns based on finite samplings of line projections.

## RESULTS

### Model Overview and Assumptions

In modeling hippocampal traveling waves, we considered wave generation and wave propagation as two separate components. As an analogy, traveling waves are viewed as packets of neural activity propagating across a brain network; wave generation is dependent on the within-block setting and wave propagation was dependent on the interblock connectivity.

In light of experimental findings (37,38), our model made several key assumptions. First, hippocampal place cells increase their place field sizes along the dorsoventral axis. Second, hippocampal excitatory place cells are dominantly driven by spatial input, whereas hippocampal interneurons are dominantly driven by theta-modulated inhibition. Third, the model consists of both sparse long-range projections between neighboring blocks and relatively denser within-block connectivity. It has generally been thought that CA3 pyramidal neurons have recurrent connections that form an autoassociative network (39), whereas CA1 cells form more limited connections with each other along the longitudinal axis (40–42). Additionally, hippocampal theta waves travel not only along the longitudinal axis but also along the CA3-CA1 direction (43). Therefore, we conceived one plausible circuit mechanism for hippocampal traveling wave generation: it was first generated in CA3 through dense recurrent connections and then propagated to CA1 through the CA3→CA1 pathway, mirroring the traveling wave in CA3.

Based on the size of dorsoventral and transverse axes of the rodent hippocampus, we divided the three-dimensional hippocampal body into 12×5 blocks (Fig. 1*A*), with each block consisting of 60 leaky integrate-and-fire (LIF) excitatory and inhibitory neurons (*Materials and Methods* and *SI Appendix*, Table S1). We first set the ratio of hippocampal excitatory (E) to inhibitory (I) neurons within each block as 3:1, with respective E-to-E, E-to-I, I-to-E and I-to-I connectivity (Fig. 1*B*). The impact or sensitivity of the ratio will be discussed later. The excitatory neurons were place-modulated (“place cells”), with gradually increasing size of place fields along the dorsal→ventral and proximal→distal axes (Fig. 1*C*). The inhibitory neurons were mainly driven by theta modulation from the medial septum. In addition to within-block connectivity, we further assumed that there was long-range reciprocal yet asymmetric connectivity between neighboring blocks along the dorsoventral axis (Fig. 1*D*). For the ease of discussion, we used the word “feedforward” to denote the interblock connections along dorsoventral or proximal-distal axis. The long-range interblock connectivity could be either purely excitatory or purely inhibitory, and the two conditions were considered separately. We assumed that the magnitude of the excitatory spatial input and inhibitory theta oscillation drive were strongest in the dorsal segment and decreased gradually along the longitudinal axis (44,45). The results derived from these settings will be demonstrated in the following subsections.

**Figure 1.**
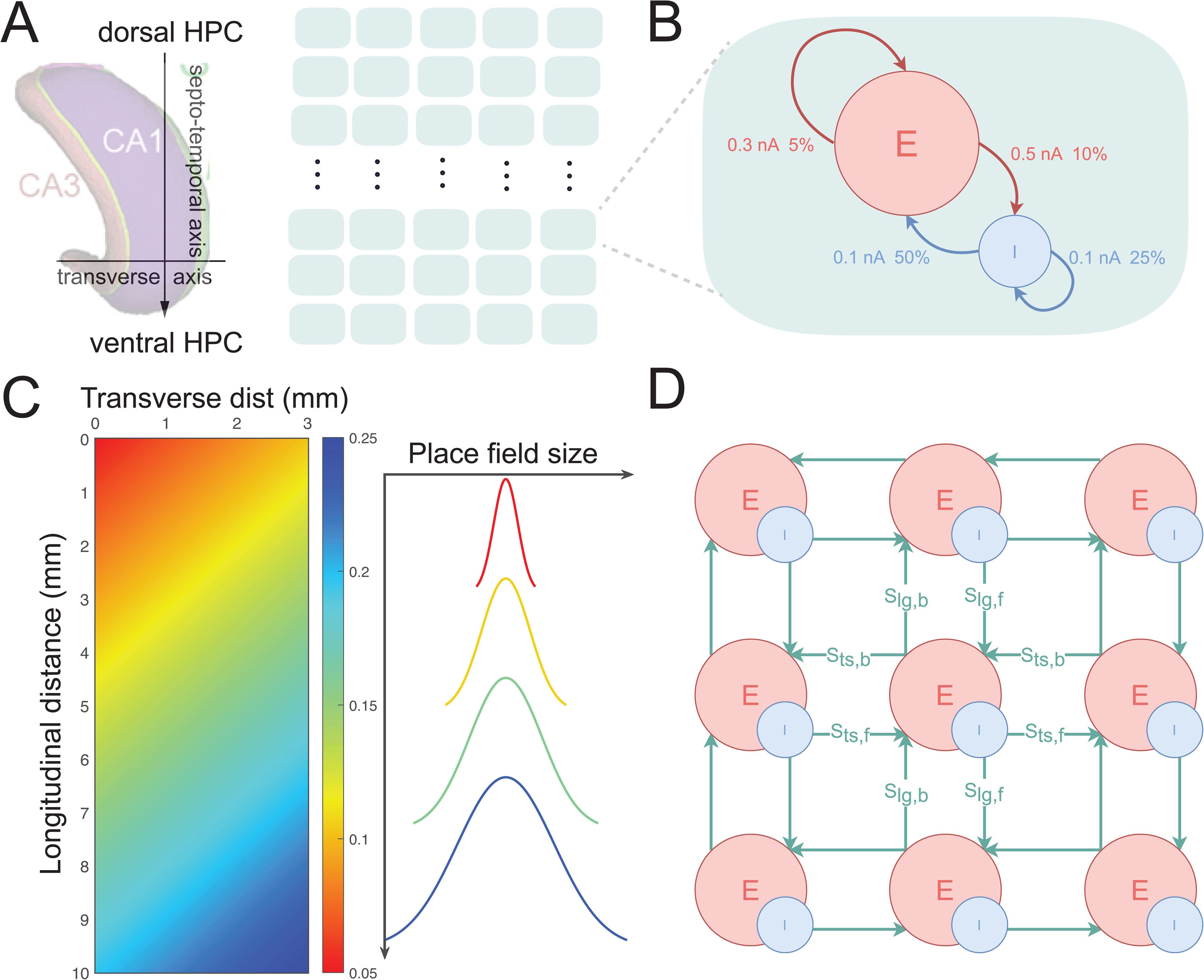
Computational models for hippocampal traveling waves. *(A)* A two-dimensional (2D) model that was mapped to hippocampal axis. The 2D model consisted of 12 blocks across the septo-temporal axis, and 5 blocks across the transverse axis. *(B)* Internal block consists of excitatory (E) and inhibitory (I) neurons. The E-E, E-I, I-E and I-E connection probabilities and strengths are illustrated. *(C) Left:* heatmap illustration of place field size in the plane. *Right:* illustration of increasing place field size along the hippocampal dorsoventral axis. *(D)* The between-block connectivity parameters {S_lg,b_, S_lg,f_, S_ts,f_, S_ts,b_}, where ***lg*** denotes longitudinal, ***ts*** denotes transverse, ***f*** denotes forward (dorsal-ventral or proximal-distal), and ***b*** denotes backward.

### Model Produces Traveling Waves That Match Findings in Rodents

Our modeling goal was to replicate the published experimental findings in the rodent hippocampus (13) (Fig. 2*A*). First, theta phase shifted monotonically with distance along the longitudinal axis, reaching ∼180° between the septal and temporal poles. Second, the power of theta oscillations decreased from dorsal to ventral sites. Third, excitatory place cells were phase-locked to the trough of local theta oscillation.

**Figure 2.**
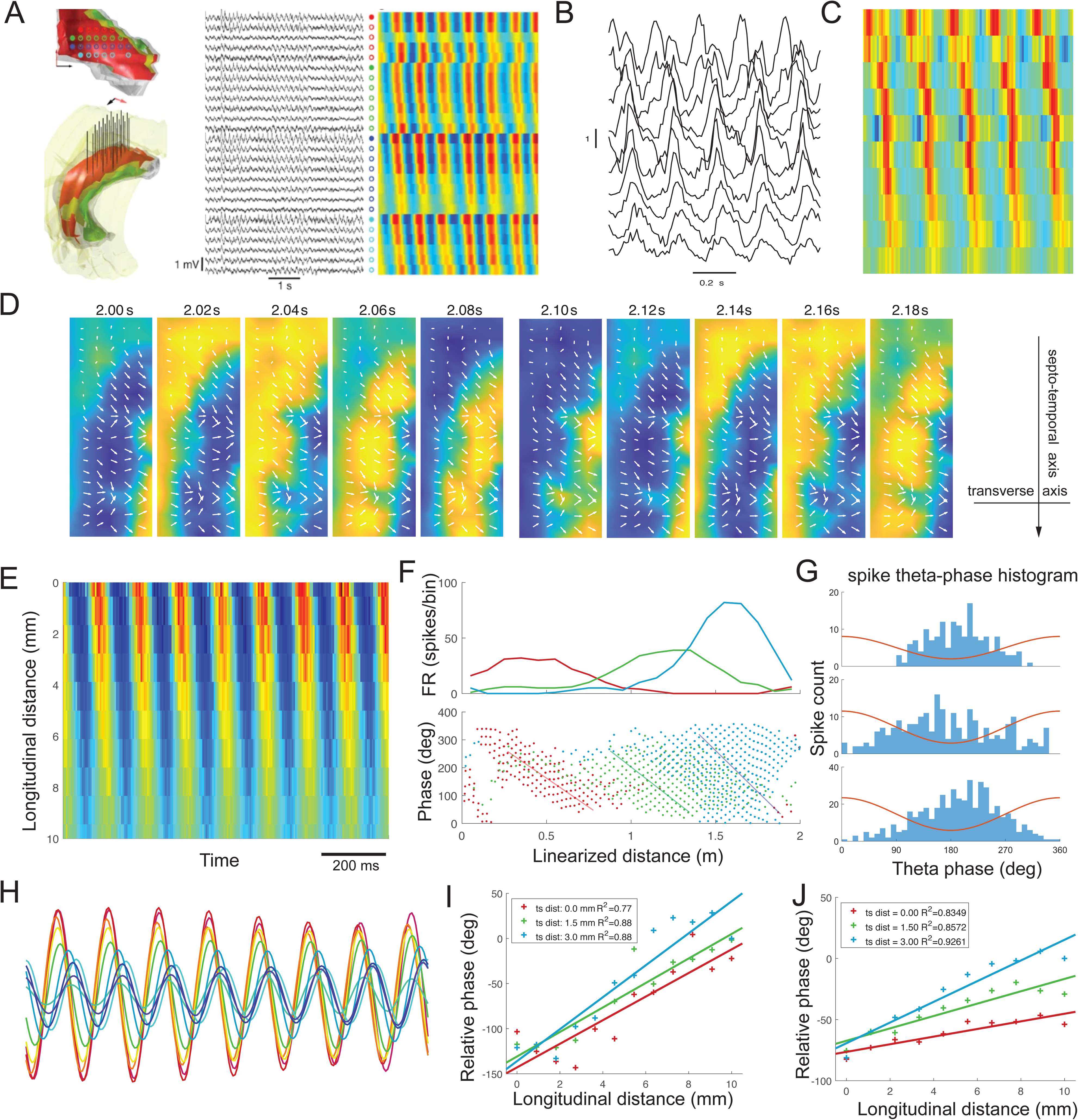
Model simulations confirm rodent hippocampal findings. *(A)* Illustration of rat hippocampal traveling waves from LFP recordings during run. *Left:* illustration of recording electrodes as a 4×8 grid. *Middle:* selected LFP traces of theta oscillations. Signals from each grid row were ordered from medial to lateral and stacked from rostral to caudal direction. *Right:* filtered LFP at theta band shown as a heatmap (Figure reproduced with permission from Springer Nature, Lubenov and Siapas, 2009). *(B)* In analogy to *A*, our model generated a proxy of 1-s LFP traces (measured as the sum of membrane potentials) in spacetime, where the activity represents the summed membrane potentials of each block (“channel”). *(C)* Heatmap representations of simulated LFP traveling wave for *B*. Note that the activity shifts along the longitudinal axis (*Top:* dorsal; *Bottom:* ventral). The activity of each channel is normalized to zero mean and unit variance. *(D)* Two-dimensional snapshots of simulated LFP traveling waves via vector field. The norm each arrow represents the gradient magnitude by log transformation for a better visualization. *(E)* Simulated block multiunit activity (MUA, by summing all spikes within the block) shifts along the longitudinal axis. Color indicates the proportion of spike activity occurring in the corresponding time period (5 ms bin). *(F)* Illustrated place receptive fields of 3 place cells (*Top*) and phase precession (*Bottom*). *(G)* Illustrated spike theta phase histograms of 3 place cells, showing spikes were fired around the trough of local theta cycle (red trace). *(H)* An illustration of 6-10 Hz filtered version of simulated LFP proxy along a dorsoventral line. Note that the theta-band activity was strongest in the dorsal segment (red) and decreased when approaching the ventral segment (blue). *(I)* Relative phase averaged over time and linear regressed to longitudinal distance. Three color lines represent the line regression at a specific distance along the transversal axis (‘ts distance’). Total phase shift of the most distal line is 177.11° (∼half cycle), corresponding to a propagation speed of 162.61 mm/s. *(J)* Similar to *I*, except for estimation from MUA.

In each simulation parameter configuration, we produced single-unit spiking activity, multi-unit activity within the block, and a proxy of local field potential (LFP) activity (which was approximated by the sum of membrane potentials in our model). At the LFP level, the model reproduced theta oscillations (Fig. 2*B*) and a spatiotemporal phase shift (Fig. 2*C*). The wave initiated at the dorsal segment (top left corner) and propagated towards the ventral segment, shifting along the transverse axis (Fig. 2*D*). We used phase gradient to illustrate the direction of wave propagation (*Material and Methods*). Similar to (12), we observed consistent traveling wave patterns at the multi-unit activity (MUA) level (Fig. 2*E*). In addition, our model produced phase precession (Fig. 2*F*), where neuronal spikes occurred progressively earlier in relation to the theta phase of LFP activity, and produced spike phase-locking phenomenon (Fig. 2*G*), where spikes were fired relative to the trough of theta cycle. From our model simulations, we observed that the proxy of theta-band LFP activity was strongest in the dorsal segment and gradually decreased towards the ventral segment (Fig. 2*H*). We characterized the phase shift vs. longitudinal distance via a linear fit (*Material and Methods*), where the phase shift slope parameter (deg/mm) was inversely proportional to the theta traveling wave speed (mm/s). We found the total phase shift of the most distal line was 177.11° (∼half theta cycle), corresponding to a propagation speed of ∼162.6 mm/s (see Fig. 2*I* for three slope estimations across the transversal axis). The phase shift slope parameter estimated from the simulated MUA appeared undershot compared to the results obtained from the simulated LFP (Fig. 2*J*). This discrepancy was probably due to the relatively low firing rate, this sparse spiking activity could not fully reflect hyperpolarization at the sub-threshold level, so the falling phase of theta oscillation was not well captured and causes the underestimation of total phase shift. In the remainder of the paper, we will focus on the results derived from simulated LFPs.

### Network Connectivity and Synaptic Strengths Determine Traveling Wave Properties

Based on the proposed network architecture, we systematically changed the network connectivity, for both within- and between-block configurations, and assessed the traveling waves derived from different configurations.

#### Case 1: Long-range excitatory connectivity

First, we considered the long-range *excitatory* connectivity only between neighboring blocks. Specifically, we systematically varied two model parameters: {S_p_, S_w0_}, and examined the relative theta phase across ten blocks along the dorsoventral axis. The synaptic weights along the longitudinal axis were reciprocal but asymmetric, being strongly biased towards the dorsoventral direction. We identified a heatmap of parameter range that produced traveling wave according to the established criterion (Fig. 3*A*). In general, in order to generate a propagating wave with proper phase shift, there is a trade-off between model parameters S_p_ and S_w0_: a smaller value of S_w0_ would require a greater value of S_p_, and vice versa. Given a fixed S_p_, the traveling wave speed was negatively correlated with S_w0_ (Fig. 3*B*). Importantly, we observed a phase transition in this model parameter. Specifically, when S_w0_ was under a threshold (∼0.4 nA in our simulations), there was no traveling wave (i.e., low magnitude of phase shift) but merely a small stable phase shift (less than 60°) induced by excitability difference. In this scenario, connections between blocks were still too weak to couple the adjacent blocks together. Since the theta activity was strongest in the dorsal segment (*Material and Methods*), each time a rising phase of oscillation occurred, the dorsal segment was first activated, followed by the intermediate and then the ventral segment. Although there was a small phase shift induced by this mechanism, it was not a traveling wave pattern since it only appeared as an epiphenomenon of the input strength difference and the activity did not propagate. As S_w0_ approached the threshold, the interaction between neighboring blocks increased and became sufficiently strong to disturb regular oscillations of independent blocks (and we called it a “phase transition”). In this scenario, the stable phase shift observed earlier on vanished, causing the R-squared metric (coefficient of determination) to decrease sharply. After S_w0_ exceeded the threshold, the interference between blocks dominated and successfully formed a chain of coupled oscillators, and the traveling wave emerged (total phase shift >60°) from such interactions. This phenomenon can be characterized in Fig. 3*C*. We also observed that the total phase shift could “overshoot” and produced a full cycle of 360° phase shift. Nevertheless, our model with optimal parameter tuning produced a phase shift of ∼180° along the longitudinal axis, as well as a phase shift of ∼40° along the transverse axis (i.e., from the subicular end to the fimbrial end of the CA1 pyramidal layer), matching the experimental findings (see Fig. S2 in Ref. 13). Furthermore, there was a gradual change in theta coherence along the longitudinal axis (*SI Appendix*, Fig. S1). We also changed the ratio of excitatory to inhibitory units (from 3:1 to 4:1, 5:1 and 9:1) and tested the sensitivity of result with respect to the overall E/I ratio (Fig. 3*D*). Generally, a higher E/I ratio affected the stability of traveling waves and caused a bit overshoot in in the phase shift (∼200°), but the overall traveling wave patterns were preserved with optimized model hyperparameters.

**Figure 3.**
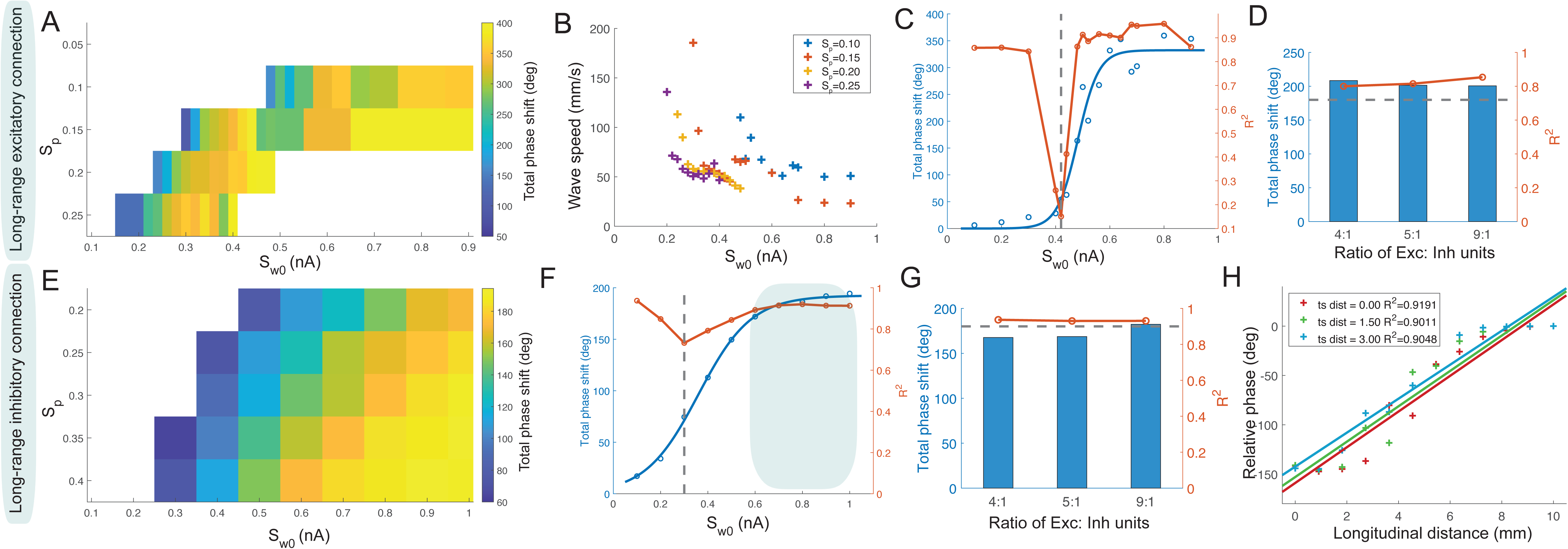
Network connectivity determines traveling wave properties. *(A)* In the long-range excitatory connectivity setup, the model parameters {S_p_, S_w0_} that produces traveling wave, where the color indicates the degree of phase shift. Conditions with R^2^ less 0.6 or phase shift less than 60° are shown in white (i.e., no stable wave propagation). Notably, ∼180° phase shift only held with constraints; if interblock connections were too dense, this phase shift could reach a full cycle. *(B)* Relationship of traveling wave speed with respect to model parameters {S_p_, S_w0_}. There is a trade-off between S_p_ and S_w0_: a greater value of S_w0_ is required for a lower value of S_p_ to attain the same speed level. *(C)* With a constant S_p_, the model went through three stages as S_w0_ increased. When S_w0_ was under the threshold (∼0.4), instead of propagating wave, there was only a small stable phase shift (less than 60°) induced by excitability difference. As S_w0_ approaching the threshold, a phase transition arose and R^c^ decreased sharply. After S_w0_ exceeded the threshold, the traveling wave emerged so that phase shift increased dramatically, and R^c^ rose again. The phase shift curve was fitted with a sigmoid function. *(D)* We altered the ratio of excitatory neurons to inhibitory neurons, while keeping the total number of neurons in each block the same (60 per block). It is noted that the total phase shift was around 200°, where the horizontal gray dash line indicates half-cycle (180°). *(E)* Similar to *A*, except for the long-range inhibitory connectivity setup. *(F)* Similar to *C*, except for the long-range inhibitory connectivity setup. Unlike panel *C*, the total phase shift was stable at 180° (shown by shaded area). *(G)* Similar to *D*, except for the long-range inhibitory connectivity setup. It is noted that the total phase shift and the corresponding **R^2^** were largely unchanged. *(H)* Under an isotropic connectivity setup, the traveling wave was prominent along the dorsoventral axis but not along the proximo-distal axis.

#### Case 2: Long-range inhibitory connectivity

Next, we considered the long-range *inhibitory* connectivity only between neighboring blocks. Surprisingly, the network produced much more stable traveling waves than the network with only long-range excitatory connectivity (*SI Appendix*, Movie S1). It also requested weaker constraints as the total phase shift did not exceed a half cycle or did not reach a full cycle even with stronger inhibitory connections. Therefore, there was a broader range of model parameters {S_p_, S_w0_} that could produce traveling wave patterns (Fig. 3*E*). Importantly, the phase shift was stable on 180° phase shift. Unlike the case of long-range excitatory connections (Case 1), it did not overshoot under a similar connection density parameter S_w0_. Additionally, the goodness-of-fit metric was better, indicating a uniform gradient along the longitudinal axis (Fig. 3*F*). Furthermore, we investigated the robustness of traveling waves in Case 2. Surprisingly, the network generated robust traveling wave patterns under a wide range of connectivity configurations (*SI Appendix,* Fig. S2 and Table S2). Additionally, the traveling wave patterns were robust with respect to the ratio of excitatory-to-inhibitory neurons (Fig. 3*G*). In the special case with isotropic reciprocal long-range connections (namely, connection probabilities and strengths of four directions were approximately the same, see Fig. 1*D*), suggesting that the anatomical constraint was not a necessary condition for wave propagation in Case 2. However, in the special case of connectivity isotropy, we only observed phase shifts along the dorsoventral axis, but not the proximal-distal axis (Fig. 3*H*).

Together, these results suggest that the network with only long-range inhibitory connectivity is flexible (in terms of reciprocal connectivity patterns) and stable for producing traveling waves. The biological plausibility of Case 1 and Case 2 models will be discussed in Discussion.

### Delay Across Theta Frequencies Supports Weakly Coupled-Oscillator Mechanism

There are two major hypotheses on the mechanism of traveling waves: weakly coupled-oscillators and fixed-delay propagation (25,46). To distinguish these two hypotheses, we calculated the absolute delay (i.e., time latency per unit distance) and relative delay (i.e., proportion of theta cycle delayed per unit distance) (*Materials and Methods*). For fixed-delay propagation models, the absolute delay should remain constant, causing the relative delay to increase as a function of theta frequency. On the contrary, for weakly coupled-oscillators models, the relative delay should not vary across different frequencies, so the absolute delay should reduce with increasing theta frequency. Consistent with experimental findings (36), our results produced a decreasing trend in absolute delay when the frequency of theta modulation input increased from 7.5 to 8.5 Hz, while the relative delay remained more stable (Fig. 4*A* and *B*), therefore supporting the weakly coupled-oscillator hypothesis.

**Figure 4.**
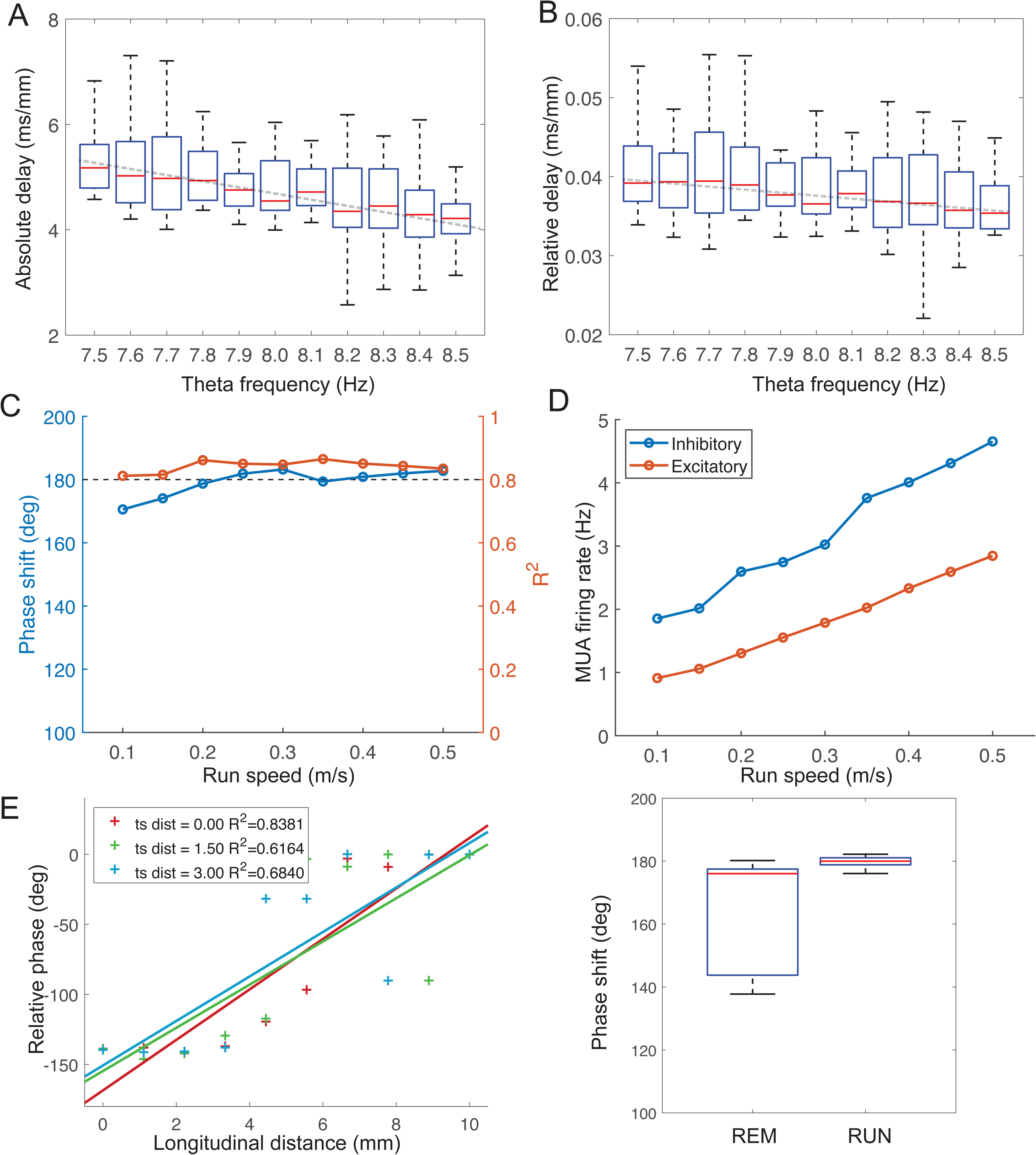
Computational model’s traveling wave properties with respect to theta frequency and speed modulation. *(A)* There was a decreasing trend in absolute delay when the frequency of theta modulation input increased from 7.5 Hz to 8.5 Hz. *(B)* In contrast, the relative delay was more stable with respect to the theta frequency modulation, supporting the weakly coupled-oscillator hypothesis. *(C)* There was a slow increase trend in total phase shift with respect to speed modulation, but the R^2^ statistics were stable. *(D)* The multi-unit activity (MUA) firing rates across all excitatory and inhibitory units increased with respect to the running speed. *(E)* In a simulated REM sleep experiment, only 15% hippocampal units were randomly activated in the model. The traveling wave patterns still persisted, with phase shift around 150°. *(F)* Box plot comparison of total phase shift statistics between REM sleep and RUN stimulations (n= 10). Each data point corresponded to the average phase shift in each Monte Carlo experiment.

### Running Speed Modulates Hippocampal Traveling Waves

Thus far, our computer simulations have assumed a default constant running speed of 0.2 m/s on the track. In the rodent literature, it is known that animal’s running speed can modulate the firing rates of place cells (47,48) as well as the amplitude of theta oscillations, especially in the dorsal hippocampus (but much less in the ventral hippocampus) (49). We further modified the Case 2-model to accommodate such running speed modulation for both hippocampal excitatory and inhibitory neurons (*Materials and Methods*). We compared the phase shift slope parameter and R^2^ with respect to running speed (0.1-0.5 m/s) (Fig. 4*C*). Additionally, our results showed that the running speed positively correlated with the multi-unit activity of both excitatory and inhibitory subpopulations (Fig. 4*D*). Notably, hippocampal theta oscillations are also prominent during REM sleep, but with two major differences in theta waves (13): slower theta frequency and lower phase shift slope (REM sleep: 16.53°/mm, total phase shift ∼150° versus RUN: 20.58°/mmm, total phase shift ∼180°). To simulate this effect, we reduced the running speed to zero in the model, and randomly selected 15% hippocampal units to be activated. As a comparison, we only randomly activated 90% hippocampal units during the RUN condition and repeated the previous analysis. We found that the traveling wave patterns still persisted in both cases, and our model prediction closely matched the experimental findings (Fig. 4*E*); the phase shift variability was also higher in REM sleep than RUN in our computer simulations (Fig. 4 *F*).

### Abnormal Hippocampal Traveling Wave Patterns

Hippocampal spatiotemporal patterns and oscillations may change according to brain states and excitation-inhibition (E/I) balance. In the pathological brains of epileptic patients, it was reported that the majority of interictal epileptiform discharges (IEDs) are indeed traveling waves that have a close relationship with seizures (50,51). In our computer simulations, we could generate “IED-like” spatiotemporal patterns based on different model parameters (Fig. 5*A*). Specifically, when the I-to-E inhibition was insufficient, traveling wave patterns still emerged initially, but the downstream blocks became over-activated so that the membrane potentials of neurons from these areas became highly synchronized and their magnitudes increased several-fold than usual (Fig. 5*B*). For instance, increasing the excitatory synaptic strengths of place cells (*I*_w0_e_, *Materials and Methods*) could produce IED-like patterns, where the magnitude of mean membrane potential could reach a large negative value (*SI Appendix,* Movie S2). Compared to the traveling waves in the normal setting, the *instantaneous* wave direction and phase shift slope had distinct patterns in the IED setting. Specifically, the wave direction distribution became less regular, compared to a Gaussian-like distribution in the normal condition (Fig. 5*C*). Furthermore, the phase shift slope parameter estimated from the IED patterns was much higher than the normal case (Fig. 5*D*). Therefore, E/I balance is critical for enabling traveling waves to propagate through the network without interrupting the original function of each local site (e.g., place selectivity). Accordingly, breaking the E/I balance may trigger the occurrence of seizures.

**Figure 5.**
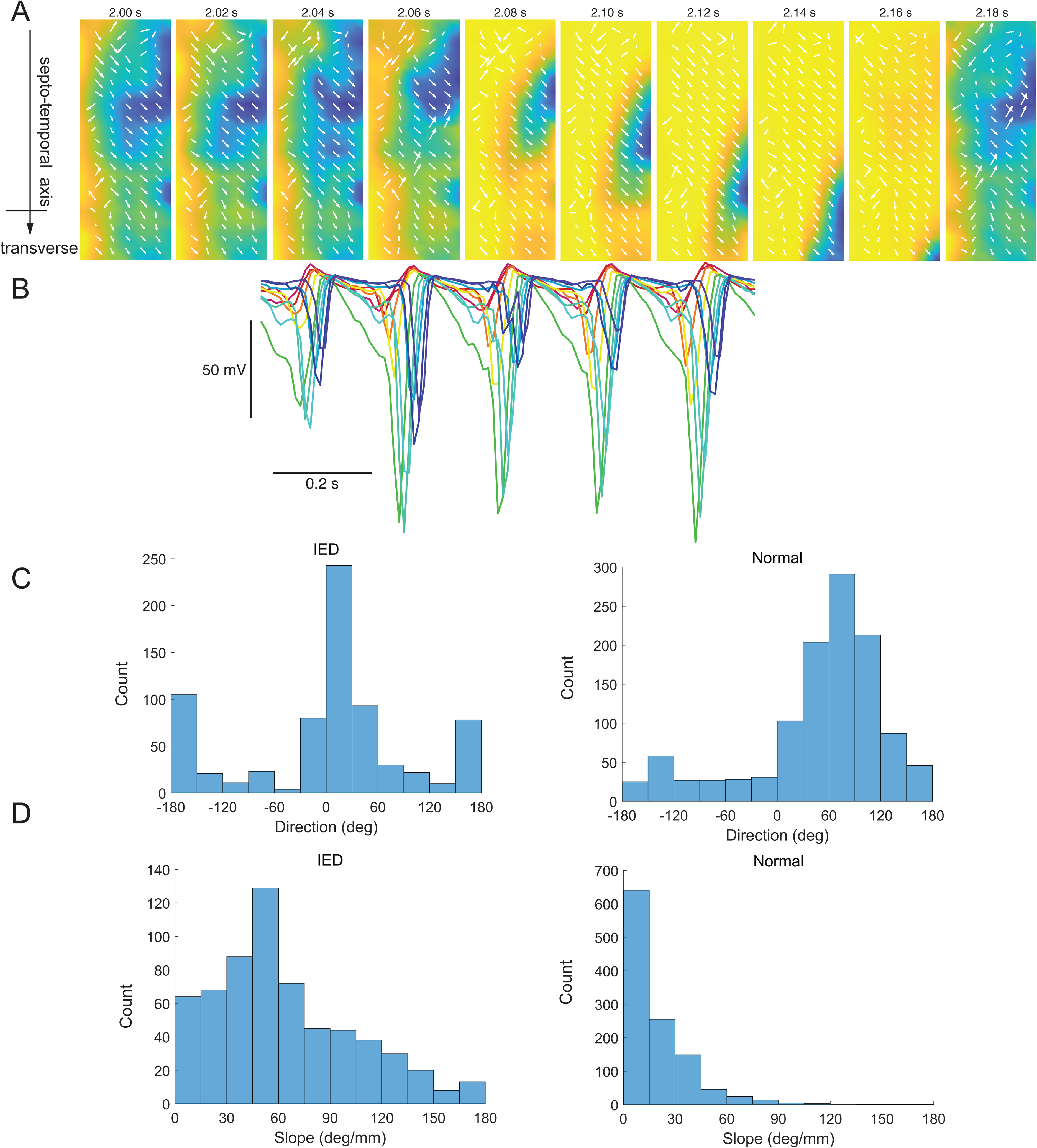
State-dependent hippocampal traveling waves. *(A)* Two-dimensional snapshots of IED-like spatiotemporal patterns. The norm each arrow represents the gradient magnitude by log transformation for a better visualization. *(B)* Simulated membrane potentials of hippocampal units became highly synchronized and substantially increased their magnitude during the IED mode. *(C)* Comparison of instantaneous traveling wave direction estimates between IED and normal conditions. *(D)* Comparison of the phase slope estimate from traveling waves between IED and normal conditions.

### Modeling Entorhinal Cortical Traveling Waves

Theta traveling waves have also recently been found in the rat MEC, propagating along the dorsoventral axis and matching the traveling waves in the hippocampus proper (36). Given long-range-projecting GABAergic bidirectional hippocampal-entorhinal inhibitory connectivity (52), one possibility is that the traveling wave was initiated in the hippocampus and then further transferred to MEC; another possibility is that the MEC generates its own independent traveling wave. To extend our model to accommodate MEC traveling waves, we changed the excitatory units in the model from 1D place cells to 1D periodic grid cells (Fig. 6*A*), while keeping other model parameters similar or unchanged (*Materials and Methods*). Similarly, grid cells show a progressive increase in grid scale along the MEC’s dorsal-to-ventral axis (53,54), and the strengths of theta oscillations decreases progressively in the dorsoventral direction of the MEC (55). Under a wide range of model assumptions, we found that our model generated traveling wave structures and properties in a similar manner (Fig. 6 *B-D* and *SI Appendix,* Movie S3), suggesting that the traveling wave patterns are robust with respect to the form of excitatory neuronal receptive fields.

**Figure 6.**
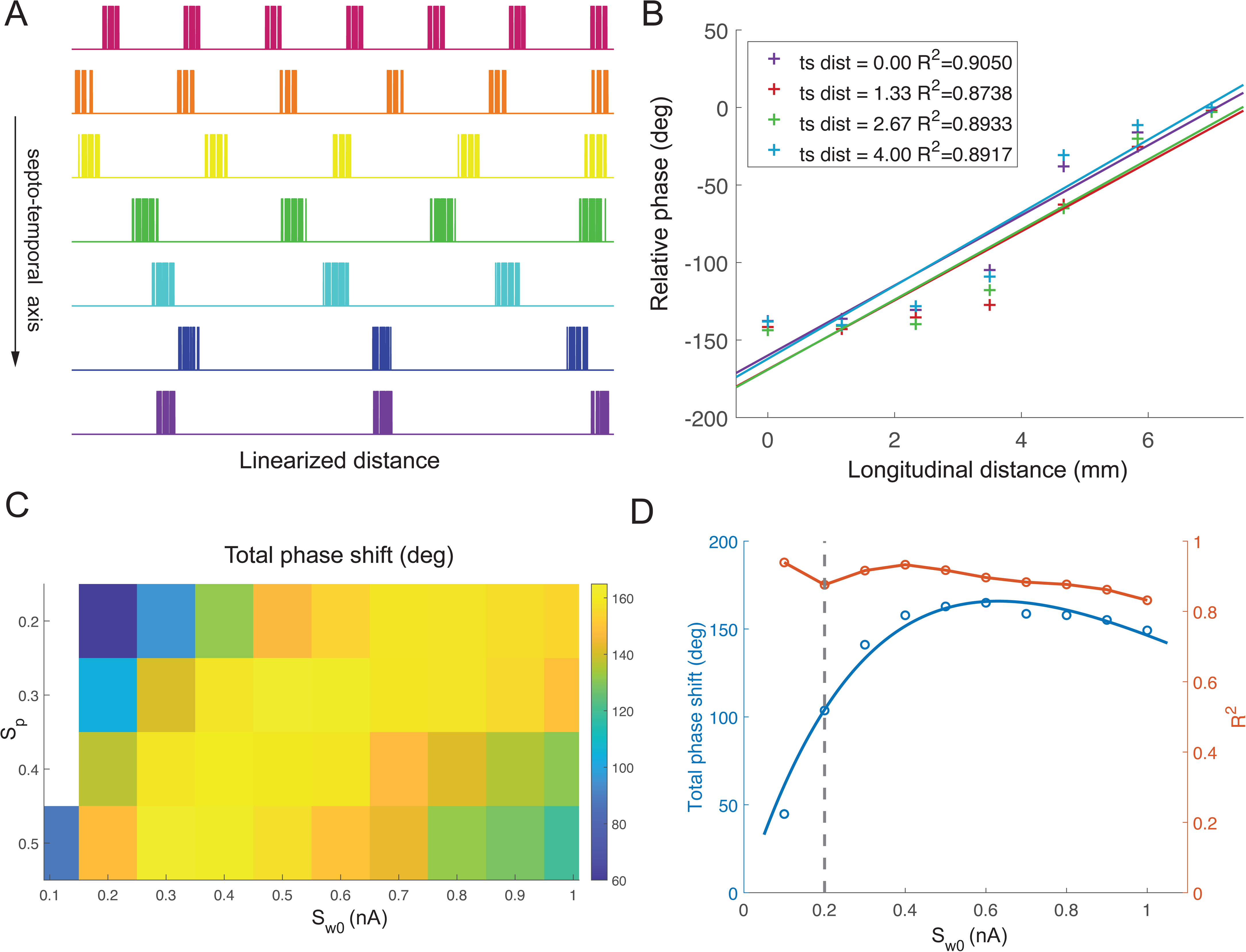
Modeling entorhinal cortical traveling waves. *(A)* Illustration of simulated MEC single-unit activity along a dorsoventral line. Each row denotes a firing of a grid unit along the 2-m linearized circular track. Note that the spacing and size of individual fields increase from the dorsal to ventral division. *(B)* Relative phase averaged over time and linear regressed to longitudinal distance. Similar to Fig. 2I, except for the MEC with grid cells. Max phase shift was 164.91°, corresponding to a slope of 23.56 deg/mm, consistent with experimental findings (26.36 ± 2.64 deg/mm, Ref. 36). *(C)* Similar to Fig. 3A, except for the MEC with grid cells. *(D)* Similar to Fig. 3C, except for the MEC with grid cells. Unlike the hippocampal case, the phase shift increased first, reached a maximum of ∼160°, and then decreased slowly. The phase shift curve was fitted with a *y* = *ax* · *e^−bx/Ä^* function.

### Detection of Traveling Wave Patterns From 1D Array Projections

Spatiotemporal traveling waves are fundamentally two-dimensional (2D) and may display unique characteristics that are often indiscernible in 1D projections. An important and practical question naturally arises: without 2D electrode array recordings, can we detect planar traveling wave patterns based on finite samplings of 1D line projections? To get further insight into this question, we first enhanced spatial resolution of our data to 51×16 by interpolation using a step of 0.2 mm. Then we performed bandpass filtering and phase estimation using the same method. To estimate phase shift slope with different projection angles, we selected 9 evenly distributed lines across a selected center (Fig. 7*A*). Since the traveling wave was more stable in the dorsal segment, we chose the center to be the dorsal-proximal point of trisection (Fig. 7*B*). For each line, we computed the relative phase as the raw phase subtracted by the phase of the most ventral-distal site on this line. We further regressed the relative phase averaged over time with respect to linear distance and computed the slope parameter and R^2^ statistic (Fig. 7*C*). Specifically, while the R^2^ metric was relatively stable, the phase slope estimate followed an “up-and-down” trend. With an increasing angle of line projection, the slope parameter increased gradually and then reached to the peak around 30°.

**Figure 7.**
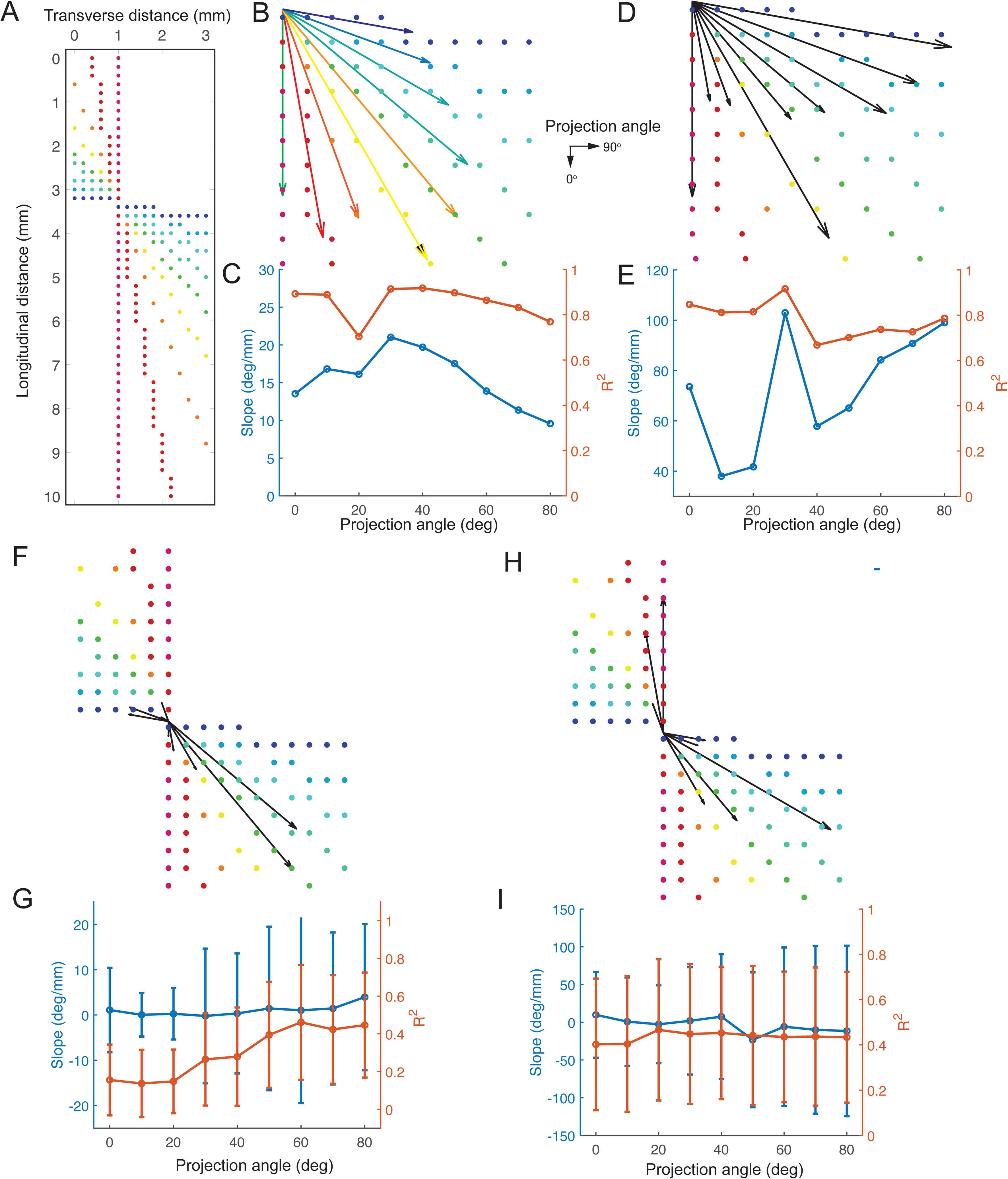
Comparison of phase shift slope based on finite samplings of line projections in two-dimensional planar traveling waves. *(A)* Illustration of sample points used to estimate the phase shift with different projection angles. *(B)* Phase shift slope along different projection lines, where a longer arrow indicates a sharper gradient. The length of each arrow is proportional to the phase shift. *(C)* The estimate of phase shift slope changes with different projection angles and corresponding R^2^ of each best-fit line, where 0° indicates dorsal-ventral axis and 90° indicates proximal-distal axis. Note that the slope reaches its local maximum at 50° and decreases on both sides. *(D,E)* Similar to *B*, except for IED patterns. Note that the estimate of IED phase shift slope changed with different projection angles and corresponding R^2^ of each best-fit line. Note that the slope was much greater than the result in *C*, and had a less regular pattern, indicating the abnormality in the spatiotemporal discharge. *(F,G)* Similar to *B*, except for spontaneous activity based on the permutation approach. *(H,I)* Similar to *B*, except for spontaneous activity based on the random noise approach.

We further repeated the same analysis on the IED-like spatiotemporal patterns (Fig. 5*A*). In this case, we observed slightly different trends in phase shift slope and R^2^ (Fig. 7 *D* and *E*). Specifically, the estimate of phase shift slope was much greater than the normal condition, and had an irregular pattern, suggesting the abnormality in spatiotemporal discharge.

As another control comparison, we generated completely random spatiotemporal patterns and repeated the same analysis. To simulate spontaneous activity, we applied both permutation and random noise approaches. In the permutation approach, we randomly reorganized the locations of all channels and used the permuted data as the input. In the random noise approach, we replaced the original signal with uniformly distributed random noise. In both approaches, we implemented interpolation, filtering and phase shift slope estimation in the same manner. We repeated these procedures 100 times and reported the mean phase shift slope and R^2^ for each line projection direction. Although the phase shifts appeared sporadically in some cases, the mean slope was close to 0 and the mean R^2^ was lower than 0.5, indicating the absence of robust spatiotemporal patterns (Fig. 7 *F-I*). Together, these results manifest a practical means to detect planar traveling wave patterns based on samplings of multiple 1D projections when implantation of dense 2D electrode arrays in deep subcortical areas is technically difficult.

## DISCUSSION

To date, many computational models have been proposed for hippocampal information processing (such as theta precession, theta and ripple sequences) (48,56–59). On the other hand, computational models of hippocampal or entorhinal cortical traveling waves have not been investigated. We developed computational models for generating mesoscopic hippocampal traveling waves. Because of the expected coupling between the entorhinal cortex and hippocampus, we also examined whether our model can be generalized to theta wave propagation in the MEC. One of key rodent experimental findings is that theta phase shifts monotonically along the longitudinal axis, reaching ∼180° phase shift between the septal and temporal poles. These results led us to reason that the half-cycle 180° phase shift is contributed by two sources: one is induced by the difference in excitability drive, and the other by propagating waves. As the dorsal and ventral hippocampal divisions have distinct anatomical output projections, the opposite theta phase across the dorsoventral axis may allow the downstream structures of hippocampus to decipher differential readout from the information carrier.

Compared to the network with only long-range excitatory connectivity (Case 1), our model simulations predicted that the network with only long-range inhibitory connectivity (Case 2) is more robust and stable, accommodating a wide range of reciprocal interblock connectivity patterns. Our computational models are based on two key assumptions: (i) there are recurrent synaptic connections within local excitatory and inhibitory neuronal populations; and (ii) there are long-range projections (either exclusively excitatory or inhibitory) along the dorsoventral axis, and these projections are reciprocal. The first assumption is justified by the well-established recurrent structure in CA3, and possibly weaker connections in CA1 (41). The second assumption is justified by the long-range GABAergic inhibitory projections in the hippocampus (60,61), and anatomically asymmetric long-range excitatory projections in CA3 may be feasible (62). Our computer simulations of hippocampal traveling waves point to a more flexible and plausible neural implementation based on long-range inhibitory connections and fewer model constraints. Our results also support the theory of coupled oscillator model (25). Recently, it has been suggested that dorsoventral traveling waves and phase precession gradients can emerge from a common theta oscillation pacemaker drive (63). One possibility for generating the dorsoventral phase shift is that there is a gradient in excitatory inputs to interneurons (or alternatively a gradient in input resistance or some intrinsic membrane current) along the dorsoventral axis. Yet another possibility is the relatively decreased fraction of inhibitory interneurons in the ventral hippocampus.

What is the impact of the lopsided connectivity pattern in the network architecture? An input gradient along the dorsoventral axis alone can induce a stable phase shift, which implies that even if both poles receive synchronous spatial inputs, the dorsal hippocampus will emit spikes earlier. Although this only accounts for a small part of total half theta-cycle delay (∼50 ms), it is sufficient to give rise to spike-timing dependent plasticity (STDP) between presynaptic and postsynaptic neurons, which will further strengthen dorsoventral-predominant connections and in turn reinforce unidirectional propagating waves. One way to test this “plasticity” hypothesis is to examine hippocampal traveling wave patterns and spatial tunings along the dorsoventral axis during the course of postnatal brain development.

What are the functional roles of hippocampal and entorhinal cortical traveling waves? In the literature, it was hypothesized that traveling waves support long-range coordination of anatomically distributed circuits, information propagation and neural plasticity (17). In addition to theta oscillations, accumulating evidence has suggested that several other brain oscillations are indeed traveling waves that may mediate information in a state-dependent manner (21,64). Bidirectional, state-dependent brain-wide interactions between the hippocampus and other cortical-subcortical areas have been observed in large-scale neural recordings (65). A plausible explanation is that computational messages occur in packages (or “chunks”) and these messages are built up in both time (66) and neuronal space. Consequently, a downstream reader interpreting the message segment should have the information about the time frame and the anatomical source of the messages. This is likely the case between the entorhinal cortex and the hippocampus. In both structures, the spatial scale of distance representations increases: place fields increase in size from dorsal to ventral hippocampus and the grid size increases from the dorsal to ventral segments of the MEC. It is also expected that these “modules” communicate with each other via theta oscillations (66). Therefore, the rules of theta propagation may be similar in both hippocampus and entorhinal cortex. Our model predictions therefore are that (i) theta oscillations also travel in the entorhinal cortex and (ii) the corresponding anatomical segments in the two structures will be in phase. With this mechanism, downstream structures of the hippocampal formation can infer the “putative position” of the animals when both theta phase and anatomical source information is available for them. This readout then should integrate with neuronal spiking sequences along the septo-temporal axis and further translate it to inferred position.

Traveling waves are expected to be an integral part of neuronal computation and messaging. Recently, it has been found in human ECoG recordings that delta (1-4 Hz) activity propagates from the cerebral cortex to the hippocampus during awake state, and the propagation direction becomes reverse during slow wave sleep, suggesting a state-dependent hippocampal-cortical dialogue (67). During working memory or rest, hippocampal theta phases can organize the reactivation of large-scale brain-wide representations (68). It is noteworthy that bidirectional propagation of low frequency (1-15 Hz) oscillations have been found on the human hippocampal surface (15). However, this finding differs from rodent hippocampal traveling waves in several aspects. First, the recording was based on electrode microgrids placed approximately on the surface of hippocampal CA1, but it did not capture the dorsoventral axis of the hippocampal CA1 or CA3 subfield. There are also anatomical differences between the rodent dorsoventral axis and the primate anterior-posterior axis in the hippocampus (4). Second, the frequency band was broader and faster than theta oscillations. Third, the subjects were engaged in a visual naming task. Independently, recent data have shown that IED or seizure activity patterns observed from the microelectrode arrays implanted in human epilepsy patients display bidirectional traveling waves, with one predominant and a second, less frequent antipodal direction (51,69). Therefore, it remains to be determined whether the bidirectionality of hippocampal or MEC traveling waves is specific to tasks or species.

In addition to the external drive, the synaptic strengths and connection probability may affect the properties of propagating waves. We hypothesize that many complex traveling wave patterns (e.g., planar, spiral, and rotation waves) reported in neocortical areas can be modeled based on a similar computational principle. Temporal delays contributed by neural transmission pathways will add additional level of complexity to modulate the frequency of theta oscillation or traveling wave patterns (31,56). It remains untested, experimentally and computationally, whether traveling waves patterns can be controlled externally to modulate the task behavior or brain state.

## MATERIALS AND METHODS

### Computational Models

#### Neurons and synapses

Both hippocampal excitatory and inhibitory populations of neurons were built based on a leaky integrate-and-fire (LIF) model. The membrane potential *V* of each neuron was determined from the following ordinary differential equations:

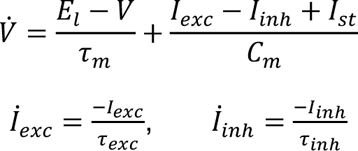

where *τ_m_* is the time constant of membrane potential, *C_m_* is the capacitance, *E_i_* is the resting membrane potential, *I_st_* is the external input current, *I_exc/inh_* and *τ_exc/inh_* denote the incoming excitatory and inhibitory synaptic currents and their corresponding time constants, respectively.

According to the dynamic differential equation, once *V* reached a predetermined threshold *V_th_* at time *t*_0_, the neuron produced a spike and then *V* was reset to the resting potential *E_i_*:

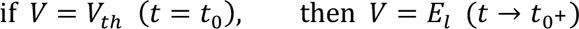

#### Excitatory place cells

We assumed that each hippocampal excitatory neuron had a specific receptive field (RF) and received a driving input containing spatial information. The external input current *I_st_* was modulated by a Gaussian-shaped function with peak located at the RF center *m_RF_*:

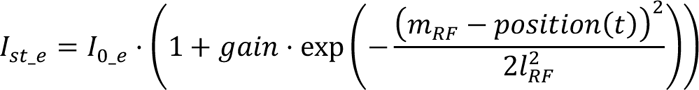

The width of this Gaussian function was determined by the RF size parameter *l_RF_*.

When an excitatory presynaptic neuron emitted a spike at time *t*_0_, a certain depolarizing current was injected to its target postsynaptic neuron after a time delay of *τ_delay_*:

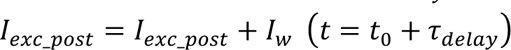

The magnitude of *I_w_* was viewed as the synaptic strength. Note that subscripts *exc/inh* denote the property of synaptic input currents (depolarizing or hyperpolarizing, respectively), *e/i* denote the excitatory/inhibitory neuronal cell type.

#### Inhibitory theta cells

The frequency of LFP theta oscillations depends primarily on the GABAergic neurons of the medial septum, and the oscillatory frequency of pyramidal neurons decreases progressively in the dorsoventral direction. We assumed that hippocampal inhibitory neurons were modulated by a pacemaker theta oscillation:

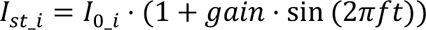

where *f* denotes the predominant frequency of theta oscillation. Here, the inhibitory neurons acted as a “hidden layer” to regulate the overall excitability of neuronal population. When a presynaptic inhibitory neuron produced one spike, a hyperpolarizing current was added to its postsynaptic neuron after a time delay of *τ_delay_*:

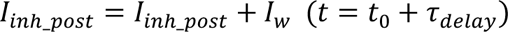

#### Local field potential (LFP) proxy

LFP consist of collective local synaptic potentials and subthreshold activity around the recording site, serving as an alternative information source about cell assemblies (70). To generate a proxy of LFP activity, we used the sum of membrane potentials of all excitatory and inhibitory cells from the model.

#### Running speed modulation

We further considered a varying running speed condition, in which the speed modulated the firing rates of excitatory place cells and inhibitory theta cells by changing the gain parameter:

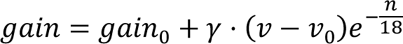

where *gain*_0_ = 1.95, *γ* = 0.8, *v*_0_ = 0.2 m/s denotes the default running speed, and *n* denotes the *n*-th block along the dorsoventral axis. The exponential decay was used to assure that there was a stronger speed modulation in the dorsal segment than the ventral segment.

#### Network setup

The whole network can be visualized in terms of a two-dimensional (2D) electrode array, with 60 recording sites altogether, namely 12 blocks along the dorsoventral (septotemporal) axis and 5 blocks along the transverse (proximal-distal) axis (Fig. 1*A*). We assumed that the excitatory place inputs ranged from 0.5 to 0.8 nA and the inhibitory theta inputs ranged from 0.4 to 0.6 nA (stronger in the dorsal segment and gradually reduced along the dorsoventral axis). The place field covered 5% to 25% of total track length (i.e., 0.1-0.5 m for a 2-m track, gradually increasing along the dorsoventral axis). Each of these parameters formed a homogeneous gradient through the 2D array.

#### Local blocks and within-block connectivity

We assumed that the hippocampus was covered by an array of local blocks in the 2D plane (or curved surface) and each block consisted of *N* neurons, which approximately corresponded to the recording scope of a single electrode. Within each block, we first assumed that 75% of units were excitatory place cells and the other 25% were inhibitory theta cells (i.e., 3:1 excitatory-to-inhibitory ratio). We also investigated the sensitivity of the ratio assumption and reported the impact of changing the ratio. It was assumed that the RF centers of place cell were evenly distributed along the whole track within each block. If the animal ran on the circuit, their place cell firing activities produced an ordered firing sequence. Based on the assumption that number of place cells was consistent over blocks, the distance between neighboring RF center was determined. Thus, the overlapping extent of neighboring RF was subject to their width (*l_RF_*). For simplicity, we further assumed that it was uniform within each block.

Synaptic connections were built within and between excitatory (E) and inhibitory (I) populations (E-E, E-I, I-E and I-I connections) with different probabilities and amplitudes (Fig. 1*B*). Additionally, we assumed that the excitatory-to-excitatory (E-E) synaptic strength between excitatory neurons decayed exponentially according to the pairwise distance of RF center with a scale factor *l_s_*:

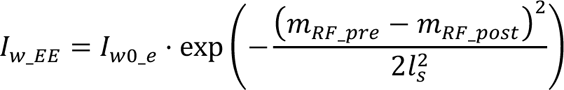

Other E-I, I-E and I-E synaptic strengths followed a normal distribution:

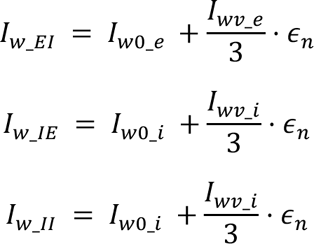

where *∈_n_* denotes a Gaussian random variable drawn from a standard normal distribution. These equations ensured that *I_w_* had a mean value of *I*_*w*0_ while fluctuating from *I*_*w*0_ − *I_wv_* to *I*_*w*0_ + *I_wv_*.

The relevant parameters were carefully selected to maintain an excitation-inhibition (E/I) balance, turning a block into a delicate oscillator which was able to alter its activity mode according to the external input.

#### Between-block connectivity

For interblock connectivity, we assumed that neurons in each block sent outgoing synapses to the adjacent blocks with certain probabilities, strengths and delays in each direction. Synapses between each block pair are denoted by a parameter group *S* = (*p_c_*, *w*_0_, *w_v_*, *t_d_*), where *p_c_* denotes probability of connection between each neuron pair, *t_d_* denotes time of synaptic transmission delay, *w*_0_ and *w_v_* determines the distribution of strengths (or weights) according to the following equation (similar to within-block connectivity):

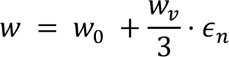

We systematically altered parameters {S_lg,b_, S_lg,f_, S_ts,f_, S_ts,b_} (Fig. 1*C*) to explore the influence of connectivity on the traveling wave features. The connection subscript *lg* denotes the longitudinal axis, *f* denotes the transverse axis, *f* denotes the forward (i.e., dorsal→ventral or proximal→distal) direction, and *b* denotes the backward direction.

##### Simulated behavior

In each simulation, we assumed that the animal ran on a circular track with a unidirectional speed *v*. The center and length of place cell RF were normalized by the total track length *l_c_*, and their values were relative (unitless). Specifically, *m_RF_* = 0 or *m_RF_* = 1 indicate that the cell fires most spikes at the end points of the linearized circular track, while *m_RF_* = 0.5 indicates that the cell fires most spikes halfway. We assumed a default running speed of 0.2 m/s on the track. For simplicity, we assumed a constant speed. A summary of model simulation parameters is shown in Table S1.

##### Model extension for entorhinal cortical traveling waves

We assumed the same 3:1 ratio for the cortical excitatory-to-inhibitory units. We further assumed the excitatory units in the MEC had grid-like tunings on the circular track (71,72). The external input current *I_st_* was modulated by a repeated Gaussian-shaped function with peaks located at its field centers (*r*_0_, *r*_0_ + *r_s_*, *r*_0_ + 2*r_s_*, …), where *r_s_* denotes the spacing between two adjacent centers, *r_0_* denotes the phase (the vertex location), and *r_n_* denotes the nearest center given the current position:

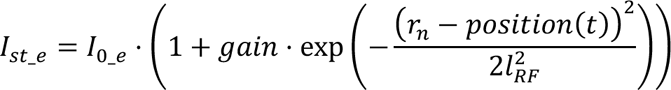

In each block, we assumed that *r*_0_ was evenly distributed along the whole track. Given that the size of MEC is smaller than the size of hippocampus (∼7 mm × 4 mm), we set 7 blocks along the longitudinal axis and 4 blocks along the transverse axis. Similarly, we assumed that three parameters formed homogeneous gradients through the 2D array (from dorsal to ventral). The excitatory grid inputs ranged from 0.8 nA to 0.6 nA and the inhibitory theta inputs ranged from 0.65 nA to 0.5 nA. The spacing between the two fields covered 15% to 40% of the total circuit length. Further, we set the field size *l_RF_* as 0.1*r_s_*. Other parameters are the same as the case of the hippocampus.

##### Software implementation

The modeling work was implemented using Brian2 (https://brian2.readthedocs.io/en/stable/), a spiking neural network simulator written in Python. The simulator is highly flexible and allows users to create non-standard neuron and synapse models. In addition, it supports Cython for speed acceleration, which can greatly improve the efficiency of computer simulation.

### Data Analysis

#### Preprocessing

The signal recorded by each “virtual electrode” was estimated by averaging the membrane potential of all *N* neurons within each block. Marginal blocks were excluded to avoid the fringe effect. The remaining channels were then filtered via a Chebyshev type-I filter with a pass band around the center oscillation frequency *f* to capture the theta band activity. This procedure greatly enhanced the stability of theta phase estimation since the Hilbert transform was only effective for a narrowband signal.

#### Instantaneous phase estimation and traveling wave characterization

For each channel, the instantaneous phase was estimated from the Hilbert transform. This absolute phase was further subtracted from the phase of the most ventral distal site (i.e., the reference node) to obtain the relative phase. A smooth filtering technique was employed in the time domain to remove the discontinuity point caused by 2π phase wrap:

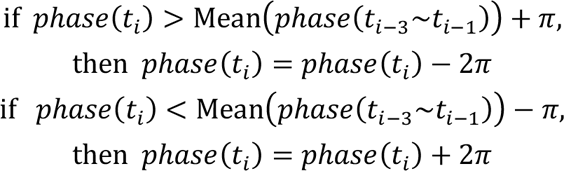

Next, this relative phase was averaged over time. The smooth filtering technique was performed again in the space domain. Namely, if the average phase difference of two adjacent blocks exceeded *π*, we performed a modulus operation (i.e., 2*π* was added to or subtracted from the phase of dorsal or proximal one since we set the most ventral distal site as the reference).

We applied a linear regression analysis between the relative phase and the relative distance (13), producing a slope parameter (unit: deg/mm). As a goodness-of-fit measure, we computed the coefficient of determination (*R*^2^) to assess the stability of traveling waves:

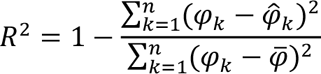

where *φ_k_* denotes the relative phase of *k*^th^ block, *φ̂_k_* denotes the phase predicted by the best-fit line, *φ̄* denotes the mean value of *φ_k_*. Thus, *R*^2^ was used to evaluate the effectiveness of linear fitting. The rationale is that for each specific direction time delay per unit distance was approximately uniform with

respect to distance. Therefore, if neural activity was traveling from one site to the other, the phase shift should be proportional to distance.

The total phase shift was defined as the product of the slope of best-fit line and the longitudinal distance. Given the phase shift *φ*, the oscillation frequency *f*, and the length of axis *d*, the propagating wave speed *s_p_* was calculated by the following formula:

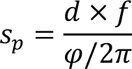

#### Visualization of traveling waves

To visualize the traveling wave, we computed the 2D vector field based on phase gradient (Fig. 2*D*). The arrow of the vector field indicates the propagation direction, and the size of the arrow is proportional to the scale.

#### Wave characterization on the cycle level

To further investigate into the change in wave property over time, we computed the cycle-by-cycle phase shift. The landmark of each cycle was defined as a peak of the theta band-filtered signal at the reference channel (i.e., the most ventral distal channel). We set the relative phase of each block to be its instantaneous phase at this landmark point, and then calculated the frequency of each cycle as the reciprocal of its duration. Based on the relative phase and theta frequency, we estimated the total shift and propagation speed on the cycle level. The relative and absolute delay of each cycle was computed as follows:

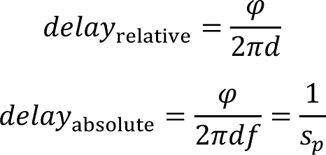

where the notations *φ*, *d*, *f* and *s_p_* represent the same parameters as on the trial level.

For each cycle, we calculated the relative phase gradient *▽φ* ≔ (*F_x_*, *F_y_*), and further estimated the propagation direction and phase shift slope of each site by arg (*F_x_* + *iF_y_*) and |*F_x_* + *iF_y_*|, respectively. Furthermore, we computed the instantaneous wave propagation speed based on the following formula:

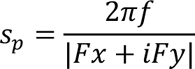

#### Analysis of multiunit spiking activity

We also tested whether multiunit activity (MUA) derived from our model exhibited traveling wave patterns. We divided the time range with a 10-ms bin size and counted the number of excitatory and inhibitory neuron spikes. We then converted MUA of each block into 100 Hz sampled time series. The relative phase, propagation speed and other parameters were estimated in the same way as the approximate LFP activity.

#### Coherence analysis

We calculated the magnitude-squared coherence between the approximated LFPs from each pair of sites using the following formula:

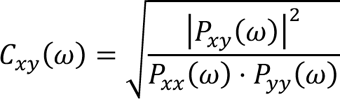

where *P_xy_*(*ω*) denotes the cross-power spectral density of *x* and *y*, *P_xx_*(*ω*) and *P_yy_*(*ω*) denote the power spectral densities of variables *x* and *y*, respectively. The theta coherence was computed by

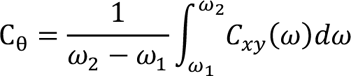

where *ω*_1_ = 4 Hz, *ω*_2_ = 10 Hz.

##### Statistics

We used the Wilcoxon signed-rank or rank-sum test for all paired and unpaired statistical tests, respectively. These tests are nonparametric and do not assume a specific distribution for the data.

##### Data and Code Availability

The data generated from computer simulations that support the findings of this study are publicly available. The code developed in this study is available on line (https://github.com/wu-yx19/TravelingWave).

## ACKNOWLEDGMENTS

We thank the G. Buzsaki for valuable comments. This work was done during a summer research internship (Y.W.). The work was supported by grants MH118928 and DA056394 (Z.S.C.) from the US National institutes of Health.

## Supporting Information for

**Figure S1:**
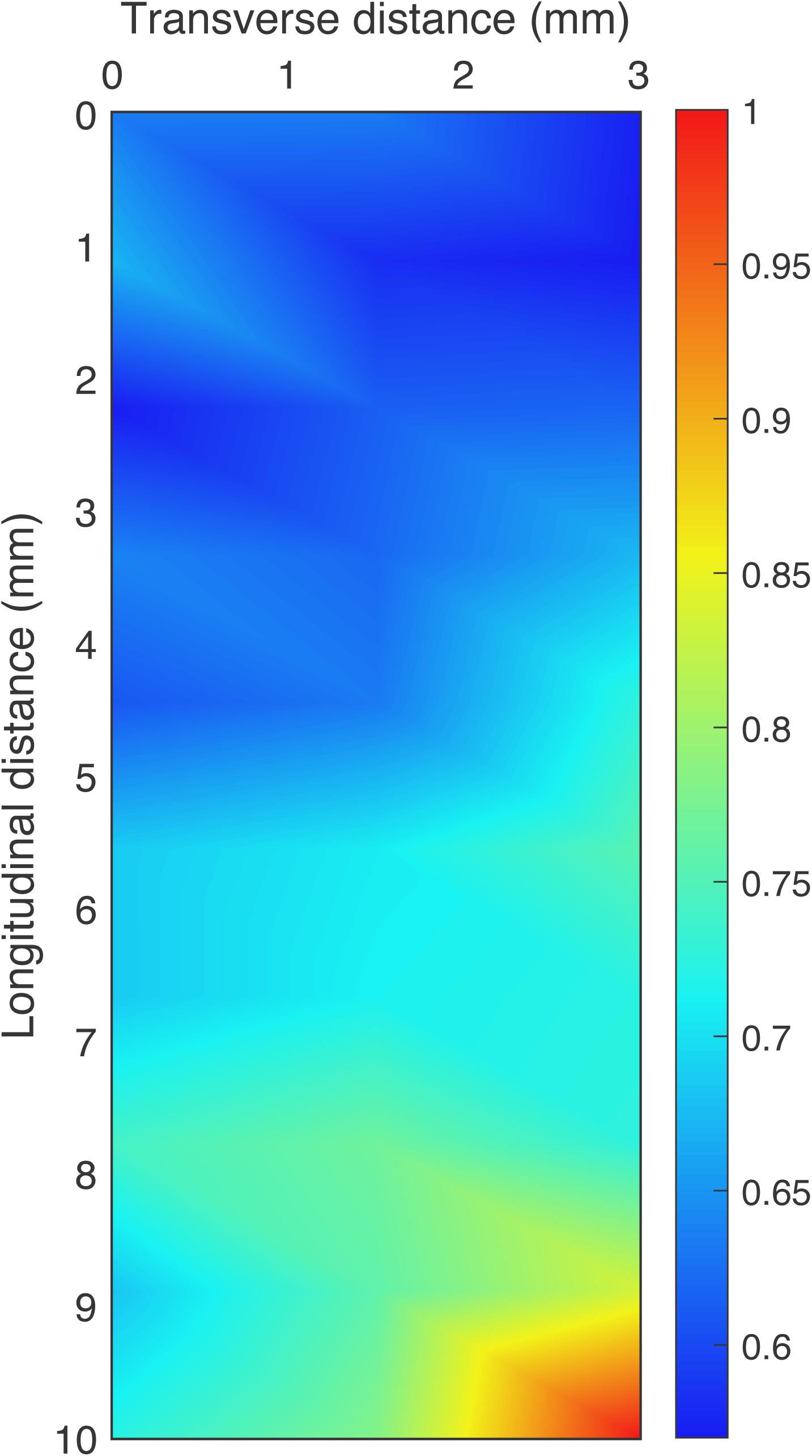
Theta coherence gradually increased along the longitudinal axis (with reference to the most ventral distal site).

**Figure S2:**
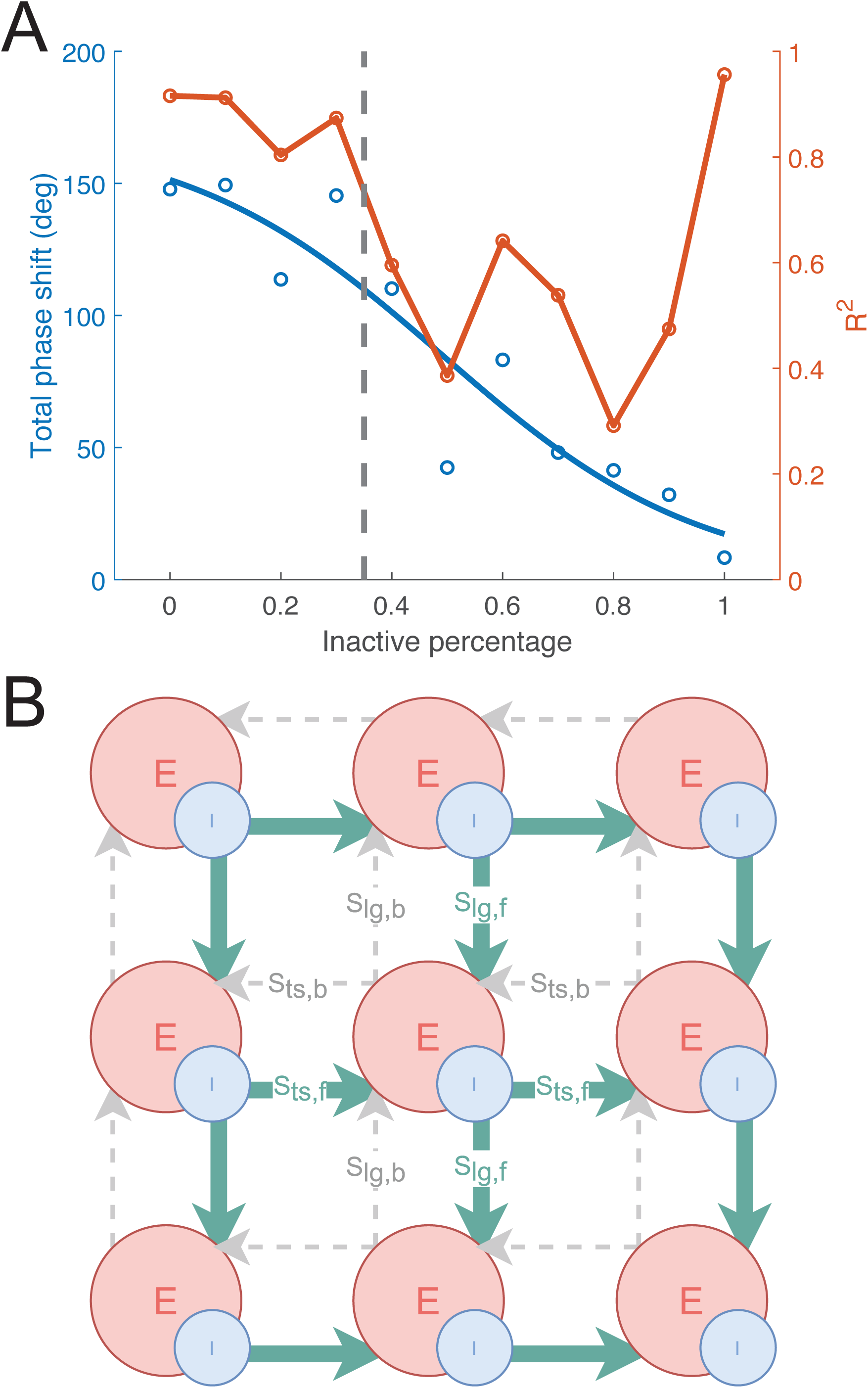
(*A*) To test the robustness of our network architecture, we randomly inactivated a percentage of long-range inhibitory synapses by setting their S_w0_ to zeros. Although the total phase shift decreased as the sparsity level increased (fitted with a sigmoid function), R^2^ remained relatively high (>0.8) when no more than 30% of synapses were inactivated, indicating that the network could still produce traveling wave patterns. Note that R^2^ became high again when all synapses were inactivated, since all recording sites were synchronized and had no phase shift. (*B*) An illustration of the “forward only” connection mode in Table S1. Gray dashed lines denote inactivated synapses.

**Table S1.**
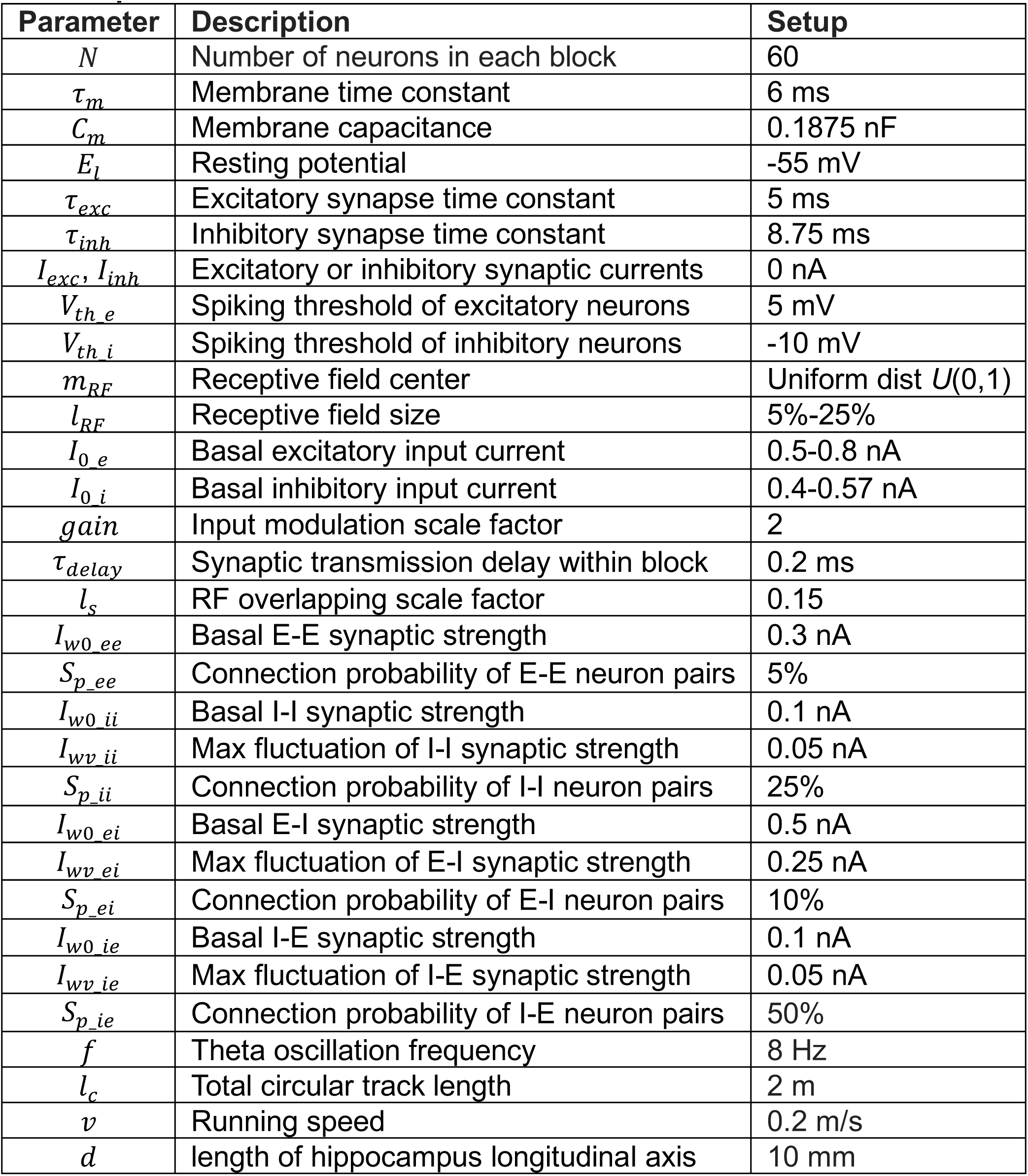
Model parameters and their default values.

| Parameter | Description | Setup |
| --- | --- | --- |
| $N$ | Number of neurons in each block | 60 |
| $\tau_m$ | Membrane time constant | 6 ms |
| $C_m$ | Membrane capacitance | 0.1875 nF |
| $E_l$ | Resting potential | -55 mV |
| $\tau_{exc}$ | Excitatory synapse time constant | 5 ms |
| $\tau_{inh}$ | Inhibitory synapse time constant | 8.75 ms |
| $I_{exc}, I_{inh}$ | Excitatory or inhibitory synaptic currents | 0 nA |
| $V_{th\ e}$ | Spiking threshold of excitatory neurons | 5 mV |
| $V_{th\ i}$ | Spiking threshold of inhibitory neurons | -10 mV |
| $m_{RF}$ | Receptive field center | Uniform dist $U(0,1)$ |
| $l_{RF}$ | Receptive field size | 5%-25% |
| $I_{0\ e}$ | Basal excitatory input current | 0.5-0.8 nA |
| $I_{0\ i}$ | Basal inhibitory input current | 0.4-0.57 nA |
| $gain$ | Input modulation scale factor | 2 |
| $\tau_{delay}$ | Synaptic transmission delay within block | 0.2 ms |
| $l_s$ | RF overlapping scale factor | 0.15 |
| $I_{w0\ ee}$ | Basal E-E synaptic strength | 0.3 nA |
| $S_{p\ ee}$ | Connection probability of E-E neuron pairs | 5% |
| $I_{w0\ ii}$ | Basal I-I synaptic strength | 0.1 nA |
| $I_{wv\ ii}$ | Max fluctuation of I-I synaptic strength | 0.05 nA |
| $S_{p\ ii}$ | Connection probability of I-I neuron pairs | 25% |
| $I_{w0\ ei}$ | Basal E-I synaptic strength | 0.5 nA |
| $I_{wv\ ei}$ | Max fluctuation of E-I synaptic strength | 0.25 nA |
| $S_{p\ ei}$ | Connection probability of E-I neuron pairs | 10% |
| $I_{w0\ ie}$ | Basal I-E synaptic strength | 0.1 nA |
| $I_{wv\ ie}$ | Max fluctuation of I-E synaptic strength | 0.05 nA |
| $S_{p\ ie}$ | Connection probability of I-E neuron pairs | 50% |
| $f$ | Theta oscillation frequency | 8 Hz |
| $l_c$ | Total circular track length | 2 m |
| $v$ | Running speed | 0.2 m/s |
| $d$ | length of hippocampus longitudinal axis | 10 mm |

**Table S2.** Qualitative results of traveling wave propagation under various long-range inhibitory connectivity configurations. Columns refer to different connection modes, where the notation “forward” denotes dorsal-ventral & proximal-distal, and “dominant” denotes that there were 50% more synapses in the dominant direction than the opposite direction. Rows refer to different levels of inactivation, where we randomly disabled a proportion of synapses to test the robustness of spiking neural network architecture. When the total phase shift was no less than 60° and R^2^ was no less than 0.7, we considered it as a traveling wave pattern and denote it with a ‘√’.

| % of inactive long-range connections | forward only | forward dominant | symmetric | backward dominant | backward only |
| --- | --- | --- | --- | --- | --- |
| 0% | √ | √ | √ | √ | √ |
| 20% | × | √ | √ | √ | × |
| 40% | × | × | √ | √ | × |
| 60% | × | × | × | × | × |

**Movie S1:** Side-by-side comparison of simulated hippocampal traveling waves for two types of long-range connectivity. *Left*: long-range excitatory connectivity only; *Right:* long-range inhibitory connectivity only.

**Movie S2:** Simulated IED spatiotemporal patterns as traveling waves. The left panel shows the run position on the circular track.

**Movie S3:** Simulated MEC traveling waves under an optimal parameter setup (S_K_ = 0.3, S_P7_ = 0.6 nA, S_PT_ = 0.48 nA, S_WX_ = 8 ms). The left panel shows the run position on the circular track.

